# Identification of regulatory elements from nascent transcription using dREG

**DOI:** 10.1101/321539

**Authors:** Zhong Wang, Tinyi Chu, Lauren A. Choate, Charles G. Danko

## Abstract

Our genomes encode a wealth of transcription initiation regions (TIRs) that can be identified by their distinctive patterns of actively elongating RNA polymerase. We previously introduced dREG to identify TIRs using PRO-seq data. Here we introduce an efficient new implementation of dREG that uses PRO-seq data to identify both uni- and bidirectionally transcribed TIRs with 70% improvements in accuracy, 3-4-fold higher resolution, and >100-fold increases in computational efficiency. Using a novel strategy to identify TIRs based on their statistical confidence reveals extensive overlap with orthogonal assays, yet also reveals thousands of additional weakly-transcribed TIRs that were not identified by H3K27ac ChIP-seq or DNase-I-hypersensitivity. Novel TIRs discovered by dREG were often associated with RNA polymerase III initiation, bound by pioneer transcription factors, or located in broad domains marked by repressive chromatin modifications. We provide a web interface to dREG that can be used by the scientific community (http://dREG.DNASequence.org).

## Introduction

Our genomes encode a wealth of distal and proximal control regions that are collectively known as transcriptional regulatory elements. These regulatory DNA sequence elements regulate gene expression by affecting the rates of a variety of necessary steps during the RNA polymerase II (Pol II) transcription cycle (Fuda et al. 2009), including chromatin accessibility, transcription initiation, and the release of Pol II from a paused state into productive elongation.

Identifying regulatory elements at a genome scale has recently become a subject of intense interest. Regulatory elements are generally identified using genome-wide molecular assays that provide indirect evidence that a particular locus is associated with regulatory activity. For example, nucleosomes tagged with post-translational modifications can be identified by chromatin immunoprecipitation and sequencing (ChIP-seq) (Barski et al. 2007; Heintzman et al. 2007). Likewise, nucleosome-free DNA can be enriched using DNase-I or Tn5 transposase (Boyle et al. 2008; Hesselberth et al. 2009; Buenrostro et al. 2013). However, each of these strategies has important limitations. Histone modification ChIP-seq has a poor resolution compared with the ~110 bp nucleosome free region that serves as the regulatory element core (Core et al. 2014; Scruggs et al. 2015; Chen et al. 2016). Likewise, nuclease accessibility assays mark a variety of nuclease accessible regions in our genomes, such as binding sites for the insulator protein CTCF or inactive regulatory elements, without the capacity to distinguish between these types of functional elements (Danko et al. 2015; Xi et al. 2007). Each of these tools is also limited by a high background, which prevents the detection of weakly active regulatory elements which may nevertheless have important functional roles.

Transcription initiation has recently emerged as an alternative mark for the location of active regulatory elements (Andersson et al. 2014a; Core et al. 2014; Danko et al. 2015). Both proximal and distal regulatory elements are associated with RNA polymerase initiation (Kim et al. 2010; Core et al. 2014; Andersson et al. 2015; Henriques et al. 2018; Mikhaylichenko et al. 2018). RNAs produced at these elements are often degraded rapidly by the nuclear exosome complex (Andersson et al. 2014b; Core et al. 2014), and as a result these patterns are most reliably detected by nascent RNA sequencing techniques that map the genome-wide location of RNA polymerase itself (Core et al. 2008; Kwak et al. 2013; Churchman and Weissman 2011; Scruggs et al. 2015). Transcription leaves a characteristic signature at these sites that can be extracted from nascent RNA sequencing data using appropriate computational tools (Melgar et al. 2011; Hah et al. 2013; Danko et al. 2015; Azofeifa and Dowell 2016).

We recently introduced dREG (Danko et al. 2015), a sensitive machine learning tool for the <u>d</u>etection of <u>R</u>egulatory <u>E</u>lements using maps of RNA polymerase derived from run-on and sequencing assays, including <u>G</u>RO-seq (Core et al. 2008), PRO-seq (Kwak et al. 2013), and ChRO-seq (Chu et al. 2018). dREG was trained to recognize characteristic signatures of nascent RNAs to accurately discover the coordinates of regulatory elements genome-wide. However, our preliminary version of dREG was limited by a slow and cumbersome implementation that made it challenging to use in practice.

Here, we present an efficient new implementation of dREG that leverages a general purpose graphical processing unit to accelerate computation. Combined with new innovations to identify regions enriched for transcription initiation (called dREG “transcription initiation regions”) and an extremely accurate new dREG model, we show that this strategy is a useful approach for detecting regulatory elements genome-wide in a number of applications. Importantly, we show that our new strategy is more sensitive in certain types of regulatory regions, for instance Pol III promoters or transcription start sites in heterochromatin domains, than DNase-I hypersensitivity or other genomic tools. Our new version of dREG is available to the community by a public web server at https://dreg.dnasequence.org/.

## Results

### A new machine learning tool for the discovery of TIRs

We recently introduced a machine learning tool for the <u>d</u>etection of <u>r</u>egulatory <u>e</u>lements using <u>G</u>RO-seq and other run-on and sequencing assays (dREG) (Danko et al. 2015). Here we introduce a new implementation of dREG which makes several important optimizations to identify regulatory elements with improved sensitivity and specificity using the multiscale feature vector introduced in dREG. We implemented dREG on a general purpose graphical processing unit (GPU) using Rgtsvm (Wang et al. 2017). Our GPU implementation decreased run-times by >100-fold, allowing analysis of datasets which took 30-40 hours using 32 threads in the CPU-based version of dREG to be run in under an hour.

We used the speed of our GPU-based implementation to train a new support vector regression (SVR) model that improved dREG accuracy. We trained dREG using 3.3 million sites obtained from five independent PRO-seq or GRO-seq experiments in K562 cells (**Supplemental Fig. S1 and Supplemental Table S1**). To improve the accuracy of dREG predictions in the unbalanced setting typical for genomic data, where negative examples greatly outnumber positive examples, dREG was trained on a dataset where bona-fide positive TREs represent just 3% of the training data. Together these improvements in the composition and size of the training set increased the area under the precision-recall curve by 70% compared with the original dREG model when evaluated on two datasets that were held-out during training (**Supplemental Fig. S2**).

We developed a novel strategy to identify regions enriched for dREG signal, which we call transcription initiation regions (TIRs), and filter these based on statistical confidence (see **Methods; Fig. 1A and Supplemental Fig. S3**). We estimate the probability that dREG scores were drawn from the negative class of sites (i.e., non-TIRs) by modelling dREG scores using the Laplace distribution. The Laplace distribution was used to model SVR scores previously (Lin and Weng 2004), and fits dREG scores in negative sites reasonably well (**Supplemental Fig. S4**). To improve our statistical power to identify bona-fide regulatory elements, we merge nearby candidate sites into non-overlapping genomic intervals, or candidate TIRs, each of which contains approximately one divergently oriented pair of paused RNA polymerases (Core et al. 2014; Scruggs et al. 2015). We compute the joint probability that five positions within each TIR are all drawn from the negative (non-regulatory element) training set using the covariance between adjacently positioned dREG scores (see **Methods**). This novel peak calling strategy provides a principled way to filter the location of TIRs based on SVR scores estimated using dREG.

**Figure 1.**
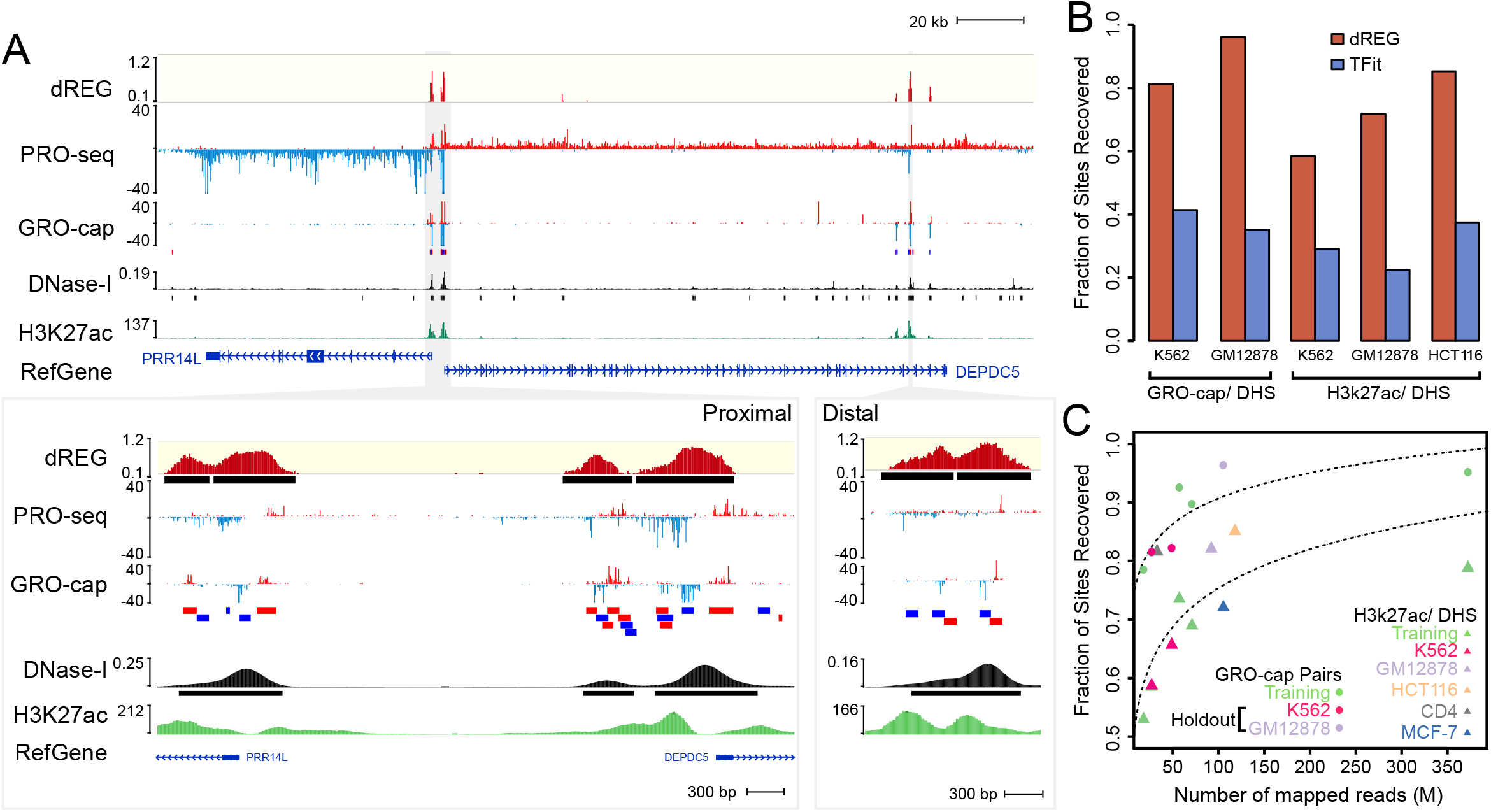
dREG identifies regions of transcription initiation. (A) WashU Epigenome Browser visualization of dREG signal, PRO-seq data, GRO-cap, DNase-I hypersensitivity, and H3K27ac ChIP-seq near the *PRR14L* and *DEPDC5* genes. Inserts show an expanded view of gene-proximal promoter elements (left) and a distal enhancer (right), each encoding multiple transcription initiation sites. (B) Barplots show the fraction of transcribed DHSs (left) and H3K27ac+ DHSs (right) that were discovered by dREG (red) and Tfit (blue) in holdout datasets. (C) Scatterplot shows the fraction of sites recovered (Y-axis) as a function of sequencing depth (X-axis) for 12 datasets shown in Supplementary Table 1. The best fit lines are shown. The color represents whether the dataset was used for training (green) or is a holdout dataset (K562, red) or cell type (GM12878, blue; HCT116, orange; CD4+ T-cells, gray; MCF-7, blue).

### Comparison to orthogonal genomic data

To evaluate the performance of dREG in real-world examples we analyzed three datasets, PRO-seq in K562 and GRO-seq in GM12878 and HCT116, that were held out during model training. Holdouts were selected because they cover a range of library sequencing depths and new cell types that together allowed us to determine whether the dREG model generalized to additional datasets. dREG predicted 34,631, 71,097, and 62,934 TIRs in K562, GM12878, and HCT116, respectively. dREG recovered the location of the majority of regulatory elements defined using orthogonal strategies at an estimated 5% false discovery rate: 81.3% or 96.1% of DNase-I hypersensitive sites (DHSs) marked by transcription (using PRO-cap pairs) and 58.4%, 71.8%, or 84.9% of DHSs marked by the acetylation of histone 3 lysine 27 (H3K27ac) (**Fig. 1B**). Sensitivity for both PRO-cap and H3K27ac-DHSs was >2-fold higher for dREG than for the elegant model-based Tfit program (Azofeifa et al. 2018) when run on the same data. Transcription initiation regions display a range in the efficiency of initiation on the two strands (Scruggs et al. 2015; Duttke et al. 2015), and dREG was able to identify the location of both uni- and bi-directional transcription initiation sites (**Supplemental Fig. S5**).

Extending dREG analysis to 14 datasets in six cell types, we found that the sensitivity of dREG varied systematically by the library sequencing depth (**Fig. 1C**). dREG achieved a reasonable sensitivity on a K562 holdout dataset with 27M uniquely mapped reads (81.3% of DHSs overlapping GRO-cap pairs were recovered), and saturated the discovery of enhancers supported by ENCODE data at between 60-100M uniquely mapped reads. After accounting for sequencing depth, we did not observe any systematic difference between datasets that were held out or used during training, suggesting that dREG was not noticeably overfitting to the training data. We did not notice any systematic bias in sensitivity for either PRO-seq or GRO-seq data, for any specific cell type, or based on the lab or origin (**Fig. 1B-C, Supplemental Fig. S6A-B**). Finally, we also obtained reasonably good performance using dREG to analyze publicly available mNET-seq data in HeLa cells (Nojima et al. 2015; Mayer et al. 2015) (**Supplemental Fig. S6A,C**). These results suggest that our new dREG model is highly extensible to nascent transcription data from a variety of different sources.

Despite a high degree of overlap with histone modification ChIP-seq assays, dREG had a higher resolution for the regulatory element core region, consisting of divergently opposing RNA polymerase initiation sites (Core et al. 2014). Regions identified by dREG were on average 6.4-fold shorter (460 bp for dREG sites) than H3K27ac ChIP-seq peaks (2,924 bp on average), closer in size to high-resolution DNase-I-seq data (322 bp on average) (**Fig. 2A**). dREG frequently separated out individual TIRs in clusters of initiation sites that could not be distinguished based on histone modification ChIP-seq peak calls, for instance in the MYC enhancer locus (**Fig. 2B**) (Fulco et al. 2016). Histone modification ChIP-seq or DNase-I-seq data aligned to the center of human dREG sites revealed good agreement with the center of the nucleosome free region (**Fig. 2C**). Thus our new dREG implementation substantially improved both resolution and accuracy compared with alternative genomic tools.

**Figure 2.**
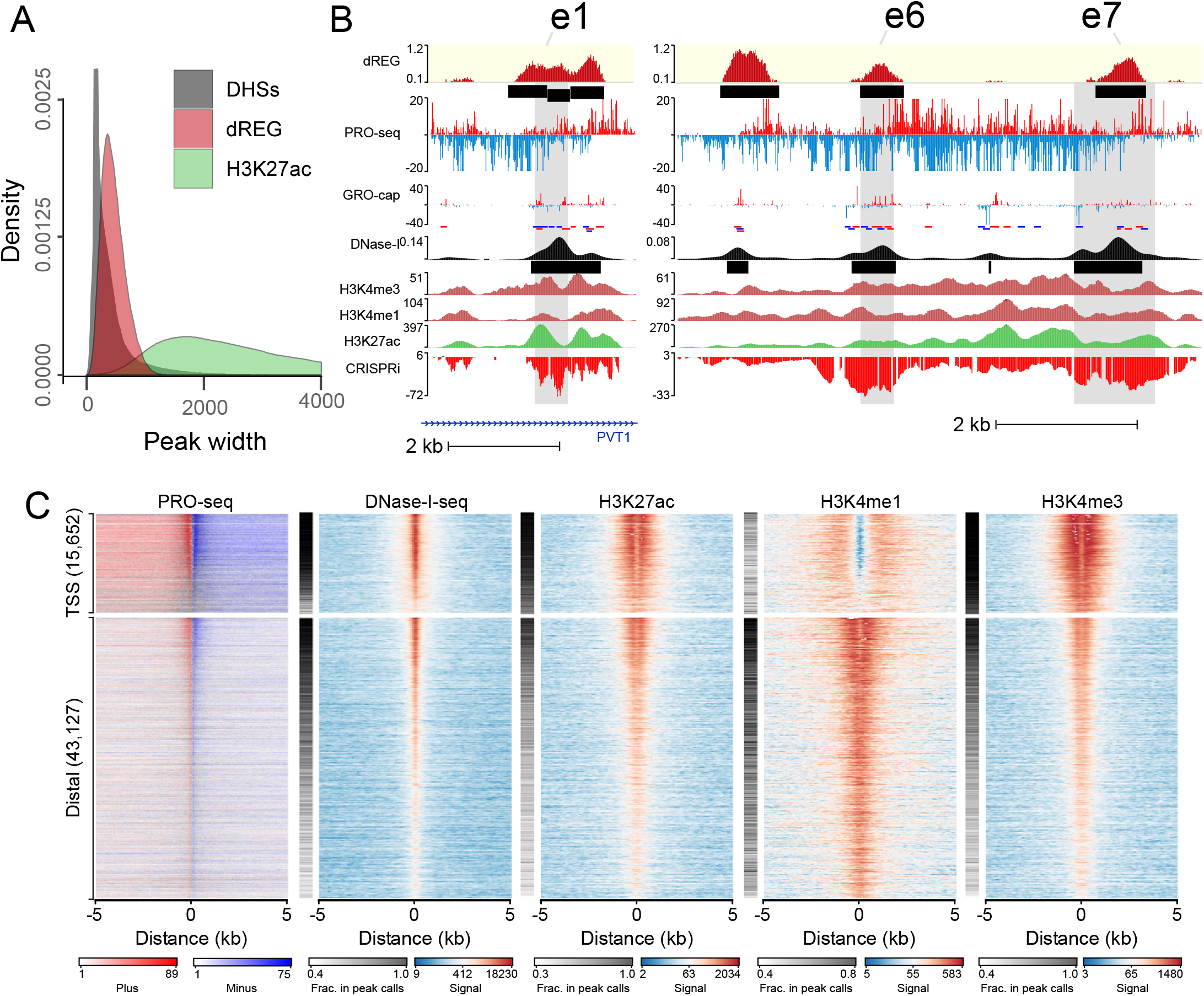
dREG calls are often concordant with other molecular assays. (A) Histogram shows the size distribution of dREG TIRs, H3K27ac ChIP-seq peaks, or DHSs. (B) WashU Epigenome Browser visualization of dREG signal, PRO-seq data, GRO-cap, DNase-I hypersensitivity, H3K4me3, H3K4me1, H3K27ac ChIP-seq and CRISPR interference score (CRISPRi) at three enhancers (e1, e2, and e3) that affect transcription of MYC in K562 cells based on CRISPR interference (CRISPRi). (C) Heatmaps show the log-signal intensity of PRO-seq, DNase-I-seq, or ChIP-seq for H3K27ac, H3K4me1, and H3K4me3. The fraction of sites intersecting ENCODE peak calls is shown in the white-black color map beside each plot. Color scales for signal and the fraction in peak calls are shown below the plot. Each row represents TIRs found overlapping an annotated transcription start site (n= 15,652) or >5kb to a start site (n= 43,127)

### Discovery of novel regulatory elements using dREG

Despite a high degree of overlap, up to 10% of TIRs did not overlap other marks expected at active enhancers. The number of TIRs found uniquely by dREG depended on sequencing depth (400-8,000 TIRs, depending on the dataset), and intriguingly did not saturate even in datasets sequenced to a depth of 350M uniquely mapped reads (**Fig. 3A**). As expected, TIRs had lower dREG scores and lower polymerase abundance when they were found uniquely by dREG (**Supplemental Fig. S7**), suggesting that these sites were often either weaker regulatory elements that were more difficult for all assays to distinguish from background, or false positives.

**Figure 3.**
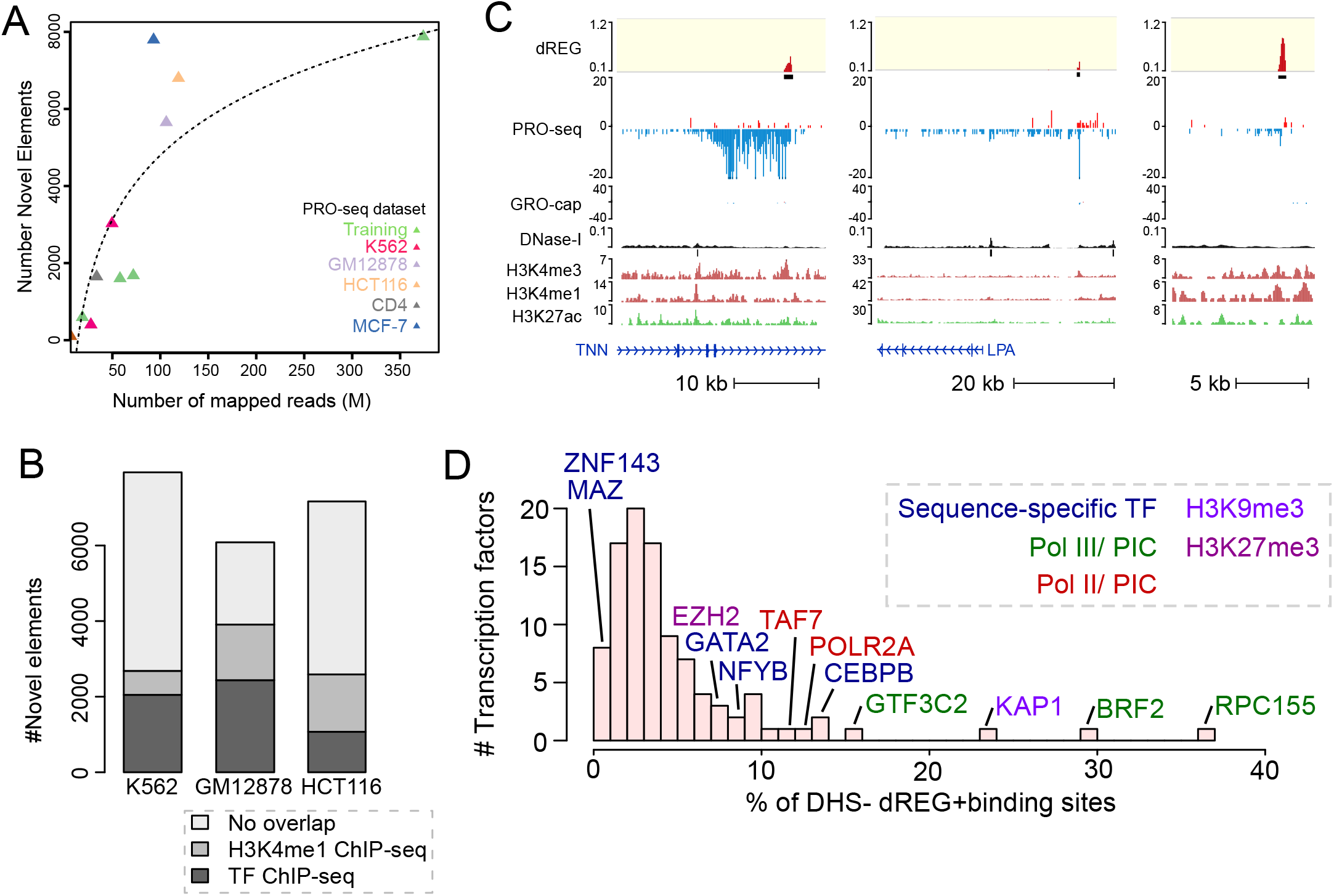
dREG identifies new regions that were not found using other molecular assays. (A) Scatterplot shows the number of new TIRs that were not discovered in DNase-I-seq or H3K27ac ChIP-seq data (Y-axis) as a function of sequencing depth (X-axis) for seven datasets shown in Supplementary Table 1. The best fit line is shown. The color represents whether the dataset was used for training (green) or is a holdout dataset (K562, red) or cell type (GM12878, blue; HCT116, orange; CD4+ T-cells, gray; MCF-7, blue). (B) Stacked barcharts show the number of elements discovered using dREG, but not found in DNase-I hypersensitivity or H3K27ac ChIP-seq (Y-axis) for PRO-seq or GRO-seq datasets in K562, GM12878, and HCT116 cells. The color denotes other functional marks intersecting sites discovered only using dREG. (C) Three separate genome-browser regions that denote TIRs discovered using dREG, but were not found in DNase-I-seq or H3K27ac ChIP-seq data. Tracks show dREG signal, PRO-seq data, GRO-cap, DNase-I hypersensitivity, H3K27ac ChIP-seq, and annotated genes. (D) Histogram representing the fraction of binding sites for 100 transcription factors supported by a dREG TIR that was not also discovered in DNase-I hypersensitivity data. Several of the outliers are shown. The color denotes whether the factor is a member of the Pol III preinitiation complex (green), Pol II preinitiation complex (red), associated with H3K9me3 (light purple) or H3K27me3 heterochromatin (purple), or is a sequence specific transcription factor (blue).

We asked whether TIRs that were not supported by DNase-I hypersensitivity or H3K27ac ChIP-seq peak calls reflect bona-fide novel regulatory elements or false positives by dREG. TIRs detected uniquely by dREG frequently (>50% depending on the dataset) overlapped ChIP-seq peak calls for sequence specific transcription factors (**Fig. 3B, Supplemental Fig. S8**). A small number of TIRs were enriched for H3K4me1, a mark associated with both active and inactive enhancers. Examining examples on the WashU Epigenome Browser (Zhou et al. 2011) revealed clearly defined transcription units that initiate long intergenic non-coding RNAs (**Fig. 3C**). Often the promoter of these transcription units lacked sufficiently robust enrichment of histone modifications or DNase-I hypersensitivity to make confident peak calls, and many lacked sufficient paused Pol II to be represented in GRO-cap data (**Fig. 3C; Supplemental Fig. S9**). Nevertheless, examination of these TIRs genome-wide revealed a local increase in the abundance of reads in the average profiles of active histone modification ChIP-seq data (**Fig. 2C and Supplemental Fig. S10**), suggesting that at least some were false negatives by other assays. Finally, sites detected only by dREG in K562 cells were often DHSs in a related cell type (**Supplemental Fig. S11**). Taken together, these findings suggest that TIRs uniquely identified by dREG were frequently novel regulatory elements, but were enriched below the level of detection of other molecular assays in K562 cells.

### Transcription factor binding predicts DHS

An alternative, but not mutually exclusive, explanation for TIRs identified uniquely by dREG is that some regulatory elements tolerate differences in the core marks reported to correlate with regulatory function. We hypothesized that certain transcription factors are more tolerant of deviations from the core regulatory architecture than others. We focused on DNase-I hypersensitivity as a general marker for the nucleosome depleted region in the center of regulatory elements. As a control for differences between K562 clones, growth conditions, or cell handling, we performed ATAC-seq to confirm low levels of chromatin accessibility in our own K562 cell stocks, closely related to those used to generate PRO-seq data (**Supplemental Fig. S12**).

To determine whether specific transcription factors may be more permissive to binding in sites having low levels of chromatin accessibility, we trained a logistic regression model to predict whether TIRs discovered using dREG intersect a DHS. Transcription factor binding site ChIP-seq data alone predicted the presence of DHSs better than using the dREG score in a matched set of holdout sites (ROC= 0.88 [TF binding], ROC= 0.75 [dREG score], **Supplemental Fig. S13**). Thus ChIP-seq data for specific transcription factors was predictive of which TIRs lacked nuclease hypersensitivity.

To identify transcription factors that contribute to this signal, we computed the ratio of ChIP-seq peak calls that were found using dREG but not DNase-I-seq, to those that were found using both assays (referred to as dREG+DHS−/dREG+DHS+). As expected, only a small fraction of most transcription factors were bound without creating a DHS (**Fig. 3D**). However, different transcription factors exhibited a broad range of binding in dREG+DHS− sites. The highest scoring outliers were frequently members of the core Pol II and Pol III transcription machinery (e.g., RPC155, BRF2, CHD1, POLR2A, and TAF7), consistent with PRO-seq detecting transcription more directly than DNase-I-seq, and potentially suggesting that some bona-fide transcription initiation sites were not sensitive to DNase-I.

### Pol III transcription initiation without chromatin accessibility

Transcription factors with the largest fraction of ChIP-seq peaks in dREG+DHS− sites were RPC155 and BRF2 (ratio of dREG+DHS−/dREG+DHS+ = 0.37 and 0.29, respectively), which encode the catalytic core of RNA Polymerase (Pol) III and a Pol III initiation factor. If a fraction of dREG+DHS− TIRs were explained by Pol III initiation, we expected to find a structured combination of DNA sequence motifs at these TIRs that were reported to be enriched in canonical Pol III promoters (James Faresse et al. 2012). Indeed, the TATA and PSE DNA sequence elements were enriched with the correct spacing in dREG+DHS− TIRs compared to TIRs that intersect POL2A (Pol II) ChIP-seq data (*p* < 1e-5; **Supplemental Fig. S14**). Enrichment of dREG+DHS− TIRs was a similar magnitude to TIRs bound by RPC155, the core subunit of Pol III, based on ChIP-seq (**Supplementary Fig. S14**). These observations suggest that some Pol III promoters were not sufficiently exposed to the DNase-I enzyme to be detected in DNase-I-seq data.

### Heterochromatin domains frequently harbor TIRs

We found that dREG+DHS− TIRs were often associated with ChIP-seq for heterochromatin markers (e.g., KAP1 and EZH2) (**Fig. 3D**). We found 6,375 dREG TIRs that overlapped heterochromatin-associated chromHMM states in K562 cells (Polycomb-repressed and Heterochromatin; low signal (Ernst et al. 2011)). In total, 55% of TIRs overlapping heterochromatin regions were not found by DNase-I-seq, a significant enrichment compared with all TIRs (*p* < 2.2e-16 Fisher’s exact test).

Next we examined TIRs in H3K27me3 domains. Broad H3K27me3 domains frequently harbored several TIRs (**Fig. 4A**). Often these TIRs were supported by GRO-cap signal, suggesting that they were not false positives. Each H3K27me3 domain contained a median of ~1 TIR per 50 kb of contiguous H3K27me3 (**Fig. 4B**). TIRs in H3K27me3 domains generally had lower levels of transcription (**Fig. 4C**), consistent with a causal role for H3K27me3 in reducing transcriptional activity (Hosogane et al. 2016; Coleman and Struhl 2017). Despite the lower levels of transcription, nearly 25% of TIRs in H3K27me3 domains were also supported by ChIP-seq for POL2A (Pol II), RPC155 (Pol III), or other transcription factors. Surprisingly, while overlap with POL2A ChIP-seq was depleted in H3K27me3 domains as expected, RPC155 ChIP-seq was over 40% enriched (*p* = 1e-8, Fisher’s exact test), potentially suggesting that Pol III initiation may be less affected by H3K27me3 than Pol II. TIRs in H3K27me3 domains also frequently overlapped active histone marks, especially H3K4me3 and H3K4me1. Taken together, our results are consistent with recent reports that transcription start sites within heterochromatin can escape repression (Leemans et al. 2018).

**Figure 4.**
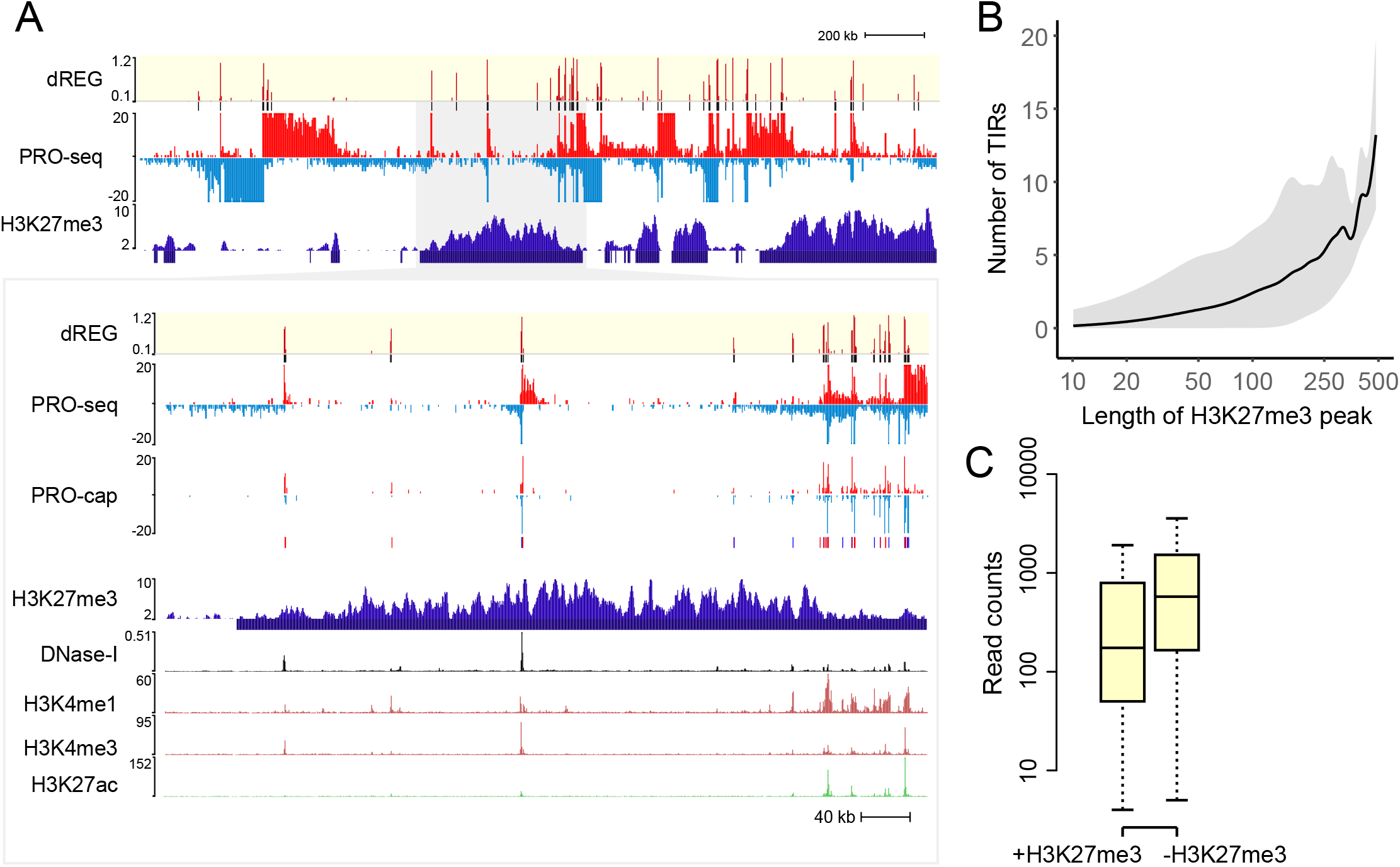
dREG TIRs located in H3K27me3 domains. (A) WashU Epigenome Browser visualization of dREG signal, PRO-seq data, GRO-cap, H3K27me3 ChIP-seq, DNase-I hypersensitivity, and H3K4me1, H3K4me3, and H3K27ac ChIP-seq. The insert shows an expanded view of the H3K27me3 domain encoding multiple transcription initiation sites that were also supported in GRO-cap data. (B) The number of TIRs discovered in each H3K27me3 broad peak as a function of H3K27me3 peak size. The line represents the median and gray shading denotes the 5th and 95th percentile. The X axis is a log scale. (C) The boxplot shows the difference in PRO-seq read counts between TIRs in an H3K27me3 peak call (+H3K27me3, left) and outside of an H3K27me3 peak call (−H3K28me3, right). The Y axis represents the number of reads found within 250 bp of each TIR.

### Transcription factors have distinct enrichment of chromatin marks in DHS− TIRs

Several sequence specific transcription factors were also observed to have a high fraction of sites that were dREG+DHS−. For example, CEBPB, NFYB, GATA2, and SPI1 had a relatively high fraction of binding sites outside of DHSs. Intriguingly, the subset of DHS+ and DHS− binding sites for these four transcription factors had distinct profiles in the flanking chromatin. All four transcription factors were enriched for increased MNase-seq read density centered on the binding site and spanning a region approximately 300 bp in DHS− sites (**Fig. 5**), suggesting systematic differences in the chromatin environment in these regions. By contrast, binding sites for MAZ and ZNF143, which exhibited a low fraction of binding sites outside of DHSs, did not show as prominent of an increase in MNase-seq signal in DHS− binding sites (**Fig 5**).

**Figure 5.**
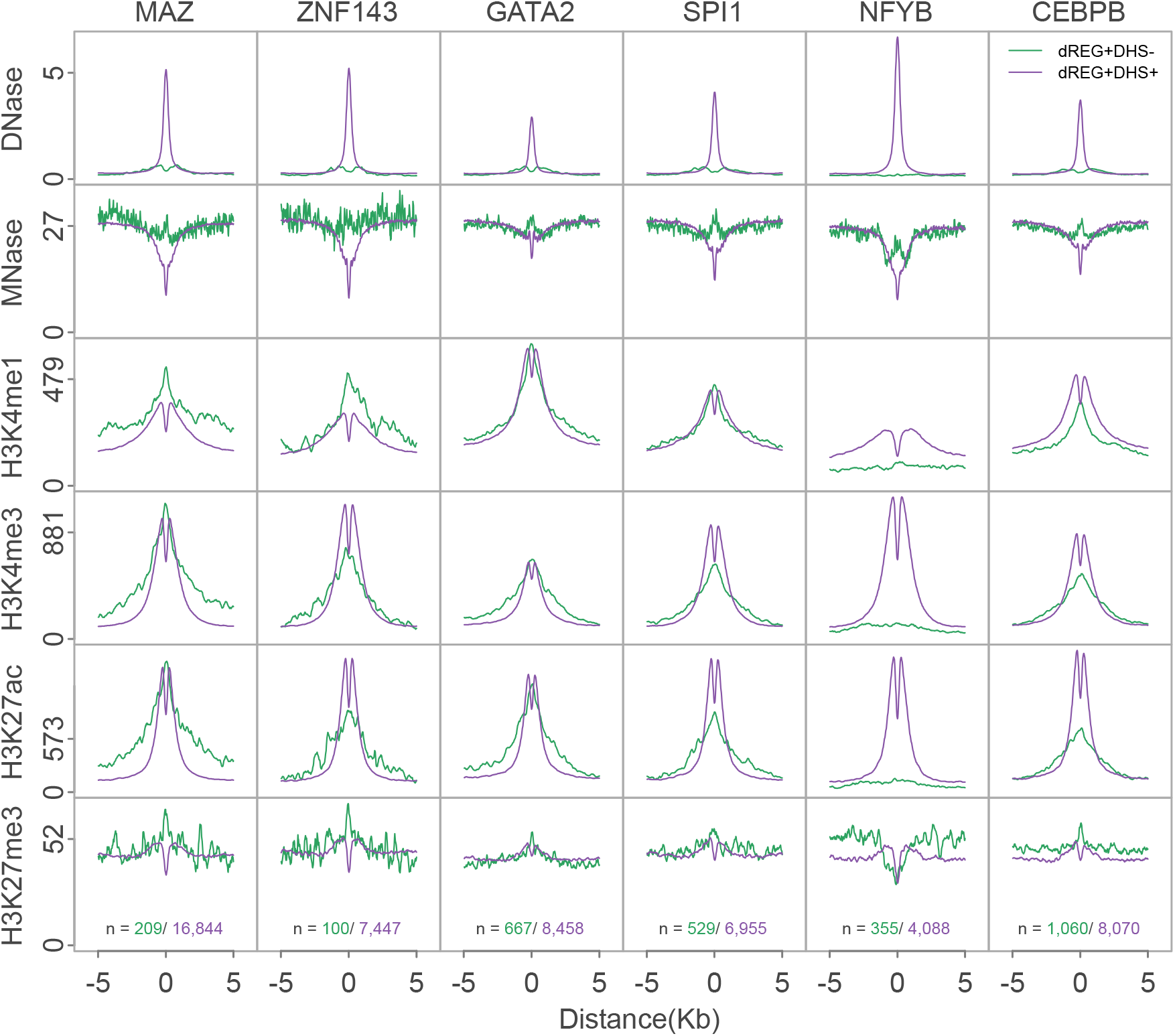
dREG TIRs with specific transcription factor binding show distinct chromatin marks. Metaplots show the raw signal of DNase-I hypersensitivity, MNase-seq, and ChIP-seq for H3K4me1, H3K4me3, H3K27ac, and H3K27me3 for six transcription factors, including MAZ, ZNF143, GATA2, SPI1, NFYB, and CEBPB. Signals are shown for dREG+DHS− (green) and dREG+DHS+ (purple) sites. The number of sites contributing to each signal is shown (bottom).

Transcription factors also showed differences in their enrichment of histone post-translational modifications. NFYB exhibited no enrichment of active histone modifications in DHS− binding sites, but was flanked on both sides by high levels of H3K27me3 (**Fig. 5**). GATA2, SPI1, and CEBPB binding sites were enriched for marks of both active and repressive chromatin, with a narrow enrichment of H3K27me3 signal localized at the putative binding site (**Fig. 5**). Likewise, histone modification ChIP-seq in DHS− regions notably lacked the dip in the center of TIRs characteristic of a nucleosome depleted region. Thus, in some cases regulatory elements discovered by dREG, but not by DNase-I-seq, appear to reflect binding of strong transcriptional activators that do not meet the current description of a regulatory element.

Taken together, these results suggest that dREG identified thousands of TIRs that were not discovered using DNase-I-seq data, but which were reproducibly associated with specific transcription factors. These observations may reflect transcription factor binding events that tolerate deviations from the core TRE architecture, preventing their discovery using more widely applied molecular tools. Collectively these observations suggest that no molecular assay has fully saturated the repertoire of active regulatory elements, even in well studied cell types like K562.

### Web server provides access to dREG

We developed a web interface for users to run dREG on their own PRO-seq, GRO-seq, or ChRO-seq data. Users upload PRO-seq data as two bigWig files representing raw counts mapped to the plus and minus strand. A typical run takes ~4-12 hours depending on server usage. Users are required to register for an account, which keeps track of previous jobs. Once dREG completes successfully, users can download dREG peak calls and raw dREG signal. Additionally the dREG web interface provides a link to visualize input PRO-seq data, dREG signal, and dREG peak calls as a private track hub on the WashU Epigenome Browser. dREG is available on the Extreme Science and Engineering Discovery Environment (XSEDE) as a science gateway (Gesing and Lawrence; Knepper et al. 2017) and is implemented using the Airavata middleware (Marru et al. 2011; Pierce et al. 2015). The dREG science gateway is available at https://dreg.dnasequence.org/.

## Discussion

In this article we have introduced an optimized version our dREG software package, a sensitive machine learning method that identifies the location of regulatory elements using data from run-on and sequencing assays, including PRO-seq, GRO-seq, and ChRO-seq (Core et al. 2008; Kwak et al. 2013; Chu et al. 2018). Our optimization efforts have achieved substantial improvements in computational efficiency, sensitivity, specificity, and site resolution. We developed a new approach to identify dREG peaks, called transcription initiation regions, based on a hypothesis testing framework that controls false discovery rates. Finally, we provide dREG as a web service where users can easily upload their own run-on and sequencing data.

Taken together, our dREG implementation has a number of advantages compared with alternative approaches. dREG offers substantial improvements in resolution for transcription factor binding sites, which tend to be located between divergently initiating RNA polymerase (Core et al. 2014). Likewise, dREG provides information about local patterns of transcription initiation, improved signal to noise ratio, and a higher sensitivity for certain types of active regulatory elements. Compared with GRO-cap (Core et al. 2014), dREG is less dependent on paused Pol II, and can also be used to detect the levels of gene transcription in the same molecular assay. Most importantly, dREG/ PRO-seq allows users to measure multiple aspects of gene regulation, including the precise position of regulatory elements, gene expression, and pausing levels using a single genomic experiment. When paired with ChRO-seq (Chu et al. 2018), which applies run-on assays in solid tissues, dREG allows the discovery of regulatory elements in primary tumors and other clinical isolates, in which the application of genomics technologies are limited by sample quantity and the cost of applying multiple assays across large cohorts.

By comparing TIRs to other functional genomic assays, we identified >8,000 regulatory elements that were not detected using DNase-I-hypersensitivity or H3K27ac ChIP-seq. Differences between assays may in part reflect false negatives in DNase-I-seq and ChIP-seq, where signals drop below the background level, or false positives by dREG. Several lines of evidence outlined in our results suggest that most TIRs are unlikely to reflect false positives. For instance, we observed a residual enrichment in the average profiles of other functional marks near TIRs that lack peak calls, which suggests that at least some fraction of TIRs reflect weak enrichment in other molecular assays that were not detected as peaks. Our results may contribute additional support to experiments assigning regulatory function to rare sites which lack canonical promoter and enhancer marks (Diao et al. 2017; Rajagopal et al. 2016). Nascent transcription may be an effective tool to expand the catalog of functional elements.

TIRs may also reflect weakly bound transcriptional activators that are relatively tolerant of binding to sites lacking DNase-I hypersensitivity. Indeed, several of the transcription factors with a relatively large fraction of dREG+DHS− binding sites were identified as having pioneer factor activity, including GATA2, SPI1, NFYB, and CEBPB (Sherwood et al. 2014; Barozzi et al. 2014; Heinz et al. 2010; Grøntved et al. 2013). It is possible that some of these elements may denote distinct architectures of functional element that are better identified using nascent transcription. Consistent with this, we found an enrichment of MNase protection at sites lacking DNase-I-seq signal. At least one of the transcription factors that we discovered having this property (GATA) was from a family reported to bind concurrently with a nucleosome *in vitro* (Takaku et al. 2016; Cirillo and Zaret 1999). Thus, one interpretation is that many of these sites reflect weak binding events in which the transcription factor and nucleosome are both present on the DNA.

A major open question following our study is whether weaker TREs that lack DNase-I hypersensitivity or other chromatin modifications have a distinct biological function. NFYB is an interesting example, as its enrichment of H3K27me3 in flanking sites, as well as a unique pattern of MNase-seq signal, might suggest binding inside of H3K27me3 chromatin domains. Transcription may be required within H3K27me3 domains either to maintain silencing, or to establish new profiles during cellular differentiation or in response to environmental signals. We anticipate that future studies will use transcription to categorize these distinct groups of functional elements in additional detail, and will determine their biological relevance in a myriad of cell types and biological conditions.

## Acknowledgements

We thank XSEDE allocation numbers TG-BIO160048 and TG-MCB160061 as well as an NVIDIA GPU Grant for providing computational resources required in this study. We thank the PHP gateway Airavata team for technical assistance building the Web service, especially Marcus A. Christie, Eroma Abeysinghe, Marlon Pierce and Suresh Marru. We also thank beta testers, Anniina Vihervaara and Donald Miller. We thank James Lewis and Anniina Vihervaara for critical comments on the manuscript, as well as other members of the Danko lab for valuable discussions and suggestions. Work in this publication was supported by an NHGRI (National Human Genome Research Institute) grant R01-HG009309 to CGD. The content is solely the responsibility of the authors and does not necessarily represent the official views of the US National Institutes of Health.

## Methods

### Overview of the dREG method

We devised a method to <u>d</u>etect the location of transcriptional <u>r</u>egulatory <u>e</u>lements from <u>G</u>RO/PRO/ChRO-seq data (dREG). The basic idea behind dREG is to differentiate between two types of regions that show high levels of RNA polymerase: (i) positions where new RNA polymerase initiates, and (ii) positions where RNA polymerase transcribes through after initiating at an upstream site. Our strategy for dREG prediction and scoring closely follows our prior work (Danko et al. 2015), except with modifications that leverage our new and considerably faster implementation to achieve higher classification accuracy. In addition, we have also added a novel strategy to improve the resolution for the region between divergently initiating transcription start sites.

We used support vector regression (SVR) to score 50 bp intervals along the genome. Loci that were low in PRO-seq reads were pre-filtered and excluded from both training and prediction tasks. We selected loci for analysis that meet either of the following heuristics: 1) contain more than 3 reads in a 100 bp interval on either strand, or 2) more than 1 reads in 1 kbp interval on both strands. These heuristics were designed to reduce the number of sites that we had to score with each dataset, while at the same time scoring at least one site near each bona-fide TIR. We refer to positions meeting these criteria as “informative positions”.

We summarized PRO-seq read counts near each position by integrating reads in non-overlapping windows centering around the informative positions, followed by transformations that are the same as in our prior work (Danko et al. 2015). Non-overlapping windows were taken at multiple scales, spanning both plus and minus strand, and both upstream and downstream directions. dREG scores can be interpreted as the degree to which each genomic position resembles a position that falls inside of a region in which transcription initiates. We use dREG scores to identify non-overlapping regions enriched for transcription initiation. We call these dREG “peaks” because they are analogous in most respects to ChIP-seq peaks.

### dREG training

The new dREG model was trained using PRO/GRO-seq signal in K562 cells obtained from five independent experiments conducted by different hands in different labs over a period of approximately two years. This diversity of training data was designed to accommodate variation in experimental conditions, batch-specific effects caused by a variety of technical factors, and detection factors such as sequencing depth. A sixth K562 dataset (G7) and a dataset representing an independent cell type (GM12878) were held out during model training to evaluate whether the final was able to generalize to additional datasets. **Table S1** lists all data sources.

PRO-seq and GRO-seq data were downloaded from Gene Expression Omnibus (see accession numbers in **Table S1**). We verified that all libraries were highly correlated with one another (**Figure S1**). Using this data, dREG was trained on a positive set of transcribed DHSs, defined as the intersection between DHSs identified by Duke and UW DNase-I-seq assays (Thurman et al. 2012), and GRO-cap HMM calls (Core et al. 2014). We defined a negative set as informative positions that do not intersect with Duke DHSs, UW DHSs, or GRO-cap HMM calls in K562 cells. We labeled each informative position as 1 or 0 according to whether it was found within a positive or negative region. To improve performance in unbalanced datasets we trained dREG on an unbalanced training set. In practice the number of informative genomic positions within and outside of bona-fide TSS differ greatly. To reduce the generalization error on genome-wide predictions, we optimized the ratio between positive and negative sets to best mimic this scenario. We selected 20K positive examples and 640K negative examples from each of the five datasets, which amounted to 3.3M training examples. Since the size of the dataset was beyond the capacity of conventional CPU-based SVM implementations, we developed a GPU-based SVM/SVR package *Rgtsvm* to handle this dataset, accomplishing the training within ~28.5hrs in a NVIDIA K80 GPU (Wang et al. 2017). The final models can be obtained from: ftp://cbsuftp.tc.cornell.edu/danko/hub/dreg.models/asvm.gdm.6.6M.20170828.rdata

### Discovering peaks enriched for dREG signal

We devised a statistical framework to identify genomic regions that are enriched for evidence of transcription initiation. We break the discovery of sites into three separate stages: First, we identify regions enriched for high dREG scores; Second, we stitch these regions into candidate peaks; Third, we estimate the probability that these peaks are drawn from the negative set of sites. Final predictions for genomic regions that contain transcription start sites are corrected using the false discovery rate correction for multiple testing and reported to the user.

During the first stage, our goal is to obtain an initial and inclusive set of sites and to stitch these into candidate peaks. We developed a statistical framework that determines a threshold dynamically for each dataset beyond which sites are likely to be located near a transcription start site. We estimate the distribution of dREG scores in negative sites using the Laplace distribution, following previous work using this distribution for the same task (Lin and Weng 2004). The Laplace distribution is parameterized by a mean and a scale (σ in **equation 1**). We assume that negative sites have a mean value of 0. The distribution of dREG scores represents a mixture distribution comprised of both negative and positive regions, and therefore fitting the scale parameter to all of the data tends to systematically overestimate the scale. To estimate the scale for a given dataset, we take advantage of the fact that the Laplace distribution is symmetric about its mean. Negative dREG scores are depleted for transcription start sites, and provide an estimate of the scale parameter which is empirically close to that obtained from the entire set of negative training examples when labels are available (**Figure S4**). Therefore, we estimate the scale parameter using negative dREG scores. Under these assumptions, the maximum likelihood estimate of the scale parameter is given as shown in (**equation 1**):

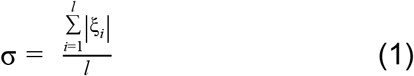

where *ξ* represents the dREG scores in training examples and *l* is the number of training examples. Genomic loci with dREG scores higher than 99.95% under the background model were selected and stitched together into intervals by extending genomic loci that pass the threshold by ±100 bp and merging these extended loci that were in 500 bp proximity. These broad regions are similar to those introduced in our first dREG publication (Danko et al. 2015).

We next designed heuristics to refine the resolution of preliminary broad regions into narrow dREG peaks. Our approach was motivated by reports that TREs often form clusters of distinct divergently oriented initiation sites within a local genomic region (Chen et al. 2016; Scruggs et al. 2015). Conceptually, our strategy increases the density of sites that are scored by dREG within the region and defines heuristics to identify local maxima. We first increased the local density of SVR predictions within the boundaries of preliminary dREG peaks, from 50 bp (in the initial prediction of broad dREG regions) to 10 bp. The dREG scores were smoothed by computing a weighted average of the seven dREG scores, representing ±60 bp of DNA (**equation 2**).

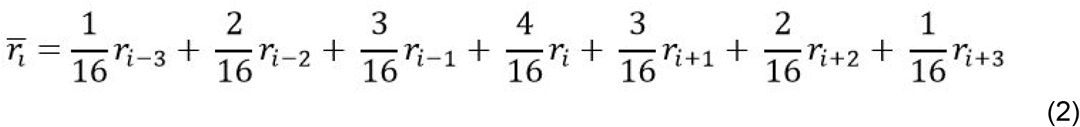

We identified points representing local maxima within each peak in which the numerical 1^st^ order derivatives changed from positive to negative. This resulted in one or more local maxima for each preliminary dREG region, each pair of which had a local minima between them. We trained a random forest to decide whether to break neighboring local maxima into separate transcription initiation regions at the local minima between them. The random forest employed dREG scores, ratio of scores between the peak and valley, and the distance each peak and the valley. The random forest was trained on a manually curated dataset on chromosome 22 of the G1 PRO-seq dataset. dREG regions that contained three or more local maxima were split iteratively until no two adjacent ignored local maxima regions existed. The boundaries of final dREG peak were defined by two valleys between the split local maxima region. For the unsymmetric broad final peaks (>= 900 bp), we trimmed the longer trail to limit the width ratio between long side and short side within 2:1. The result of this procedure was a set of non-overlapping transcription initiation regions which were often found in clusters.

To estimate the statistical confidence of each candidate dREG peak we devised a hypothesis testing framework in which we test the null hypothesis that points within each peak are drawn from the null (i.e., non-TRE) distribution. We consider five dREG scores around the peak center (i.e., peak center - 40bp, peak center - 20bp, peak center, peak center + 20bp, peak center + 40bp). Small peaks (<50 bp), were removed. We model dREG scores using a multivariate Laplace distribution parameterized by a mean vector and a covariance. We set the mean vector to 0, which corresponds to our null hypothesis that all five of these points are in negative regions. Nearby dREG scores have a complex correlation structure, requiring us to account for the covariance between sites. The covariance structure was specified by the Toeplitz matrix with homogenous variances and heterogeneous correlations (**equation 3**), because this formulation provides the most flexibility to fit complex data, and plenty of data is available for training in each dataset. We compute the variance, *σ^2^*, between sites every 20 bp using all of the dREG scores in the dataset.

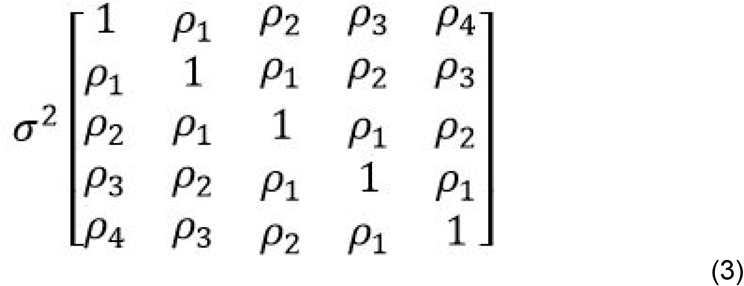

We calculated the p value based on the conditional cumulative distribution of a multivariate Laplace (i.e., p(S_i_≥ps_i_ | X_i_=0, for *i* ∈ [1,…,5]), where ps_i_ denotes the predicted score for locus *i*). Each dREG peak is associated with an estimated *p*-value. P-values are corrected for multiple testing using the Benjamini Hochberg false discovery rate (Benjamini and Hochberg 1995). By default, dREG reports peaks with an FDR corrected p value ≤ 0.05.

### Web-based implementation using Apache Arvitata

The public web-based version of dREG is hosted as a Science gateway in the Extreme Science and Engineering Discovery Environment (XSEDE) high performance computing resource (Gesing and Lawrence; Knepper et al. 2017). The dREG Gateway is hosted on the Jetstream server as a web service which can submit compute jobs and download the results of dREG peaks. From the view of software architecture, it can be divided into two parts: the secured web service and the High-performance computing (HPC) resource. The secured web service is built with PGA (PHP gateway with Airavata) on an Apache web server performs user authentication, data upload, sequence data transfer, and jobs submission to GPU servers using Apache Airavata middleware (Marru et al. 2011; Pierce et al. 2015). The HPC resources are GPU servers hosted by XSEDE. The dREG gateway, uses a job scheduler to call the *dREG* package complete the peak calling on GPU nodes. Once the calculation is completed, Apache Airavata copies the results from the HPC storage into the user’s web storage. Since this gateway uses GPUs to speed up dREG prediction with the aid of the *Rgtsvm* package (Wang et al. 2017), a typical run takes ~4-12 hours (mean = 6.7 hours) after the job starts running on the GPU server.

### Using TFit

The Tfit software (most recent on April 28th 2017) was obtained from https://github.com/azofeifa/Tfit. The Tfit software package was run using the default parameters, following instructions from the package authors. We tried using a variety of different settings (both with and without optimizing the template density function by promoter or TSS associated regions; -tss parameter). We also explored treating input data as both the full Illumina mapped read, or representing the position of RNA polymerase using the single base corresponding to the 3’ end of each read. We present the parameters that achieved the highest sensitivity for transcribed DHSs (without the -tss parameter, and using the complete Illumina read in the input bigWig).

### Comparison to DNase-I-seq and ChIP-seq data

DNase-I hypersensitive sites for the ENCODE reference cell types were processed using a uniform pipeline that we recently described (Chu et al. 2018). Sites detected using dREG were classified into DHS+ (defined as TIRs having peak calls in both Duke and UW DNase-I hypersensitivity data), and DHS- (defined as having peak calls in neither Duke nor UW data). All computations on bed files were performed using bedtools (Quinlan and Hall 2010). Bedtools was used to calculate overlap regions (*bedtools intersect*), closest distances (*bedtools closest*) and jaccard scores (*bedtools jaccard*). ENCODE hg19 blacklist regions were excluded from all analyses. Downstream processing, data analyses, heatmaps, and other and visualizations were performed in R using the bigWig package (https://github.com/andrelmartins/bigWig). Our scripts are posted on GitHub (https://github.com/Danko-Lab/dREG/tree/master/GR_submit_2018).

### Cell culture

Cell lines were obtained from the American Type Culture Collection (ATCC) and cultured using standard cell culture procedures and sterile technique. Human K562 suspension cells were cultured in RPMI-1640 media supplemented with 10% Fetal Bovine Serum (FBS) and 1% Penicillin/Streptomycin. Human HeLa adherent cells were cultured with Dulbecco’s Modified Eagle Medium media supplemented with 10% FBS and 1% Penicillin/Streptomycin. Media and antibiotics were from Corning and FBS was from Atlanta Biologicals.

### ATAC-seq data preparation and processing

ATAC-seq was performed on K562 and HeLa as described in (Buenrostro et al. 2013). Briefly, nuclei were isolated from 50,000 K562 and HeLa cells in duplicate, tagmented using the Nextera DNA Sample Preparation Kit, and amplified for 7 PCR cycles using the NEBNext DNA Library Prep Kit. All libraries were pooled and sequenced using an Illumina NextSeq500. Raw sequencing data was aligned to hg19 using bwa mem.

## Supplementary Figure Legends

**Supplemental Figure S1. PRO-seq datasets used during dREG model training**. Heatmap shows Spearman’s rank correlation (upper left) and raw gene-body correlations (lower right) between five PRO-seq and GRO-seq datasets used during dREG model training.

**Supplemental Figure S2. dREG accurately detects the location of regulatory elements**. Precision-recall curves show the precision (Y-axis; true positives/ [true positives + false positives]) and recall (X-axis; true positives / [true positives + false negatives]) of the new and previously published dREG models on the indicated dataset. The gold standard positive set was defined as DNase-I hypersensitive sites having a GRO-cap-defined transcription start site. The negative set was defined as sites lacking both GRO-cap and DNase-I hypersensitivity. The figure reflects all informative positions having a threshold of PRO-seq signal scored by dREG in the indicated dataset. Both datasets were held out during model training.

**Supplemental Figure S3. Procedure for discovering transcription initiation regions (TIRs)**. We devised a new method for finding peaks of dREG signal, called transcription initiation regions (TIRs). dREG selects informative positions and predicts dREG signal, increases the local density in high scoring regions, and smoothes to identify local increases in signal intensity.

**Supplemental Figure S4. Laplace distribution fits dREG scores in negative regions**. Histogram shows the density (Y-axis) of dREG scores (X-axis) in regions that were defined as true negatives using orthogonal sources of genomic data. The lines represent the best fits to the distribution based on all true-negative sites (blue) or based on negative dREG scores (red).

**Supplemental Figure S5. dREG identifies unidirectional TREs**. (A-C) WashU Epigenome Browser visualization of dREG signal, PRO-seq data, GRO-cap, DNase-I hypersensitivity, and H3K27ac ChIP-seq near the *ASTN1* and *HHAT* genes. TIR indicated by the gray bar (A) lacks signal in both H3K27ac and DNase-I hypersensitivity. Two TIRs (B-C) are supported by reads on only one strand. (D) Ratio of reads in a 500 bp window of the TIR center (TIR center to +/−250) on the plus and minus strand identified a large diversity in the directionality index on the plus and minus strand in the deeply sequenced K562 dataset.

**Supplemental Figure S6. The new dREG model generalizes well to other data types**. (A) Scatterplot shows the fraction of sites recovered (Y-axis) at a 5% estimated false discovery rate as a function of sequencing depth (X-axis) for 13 datasets shown in Supplementary Table 1. The best fit lines are shown. The color represents the assay that was used to obtain the data, either PRO-seq (green), GRO-seq (pink), Churchman’s mNET-seq (purple), or Proudfoot’s mNET-seq (blue). (B) Barplots show sensitivity for different datasets at an estimated 5% FDR. Sensitivity is shown for the standard dREG model (red), an alternative dREG model trained on an additional K562 GRO-seq dataset generated outside of Cornell (blue), or a dataset trained on both K562 and GM12878 (green). (C) Table shows the number of TIRs recovered, the true positive rate (TPR), and the upper-bound false discovery rate (FDR) derived using the fraction of TIRs that do not intersect peak calls in H3K27ac or DNase-I-seq data. Proudfoot data was collected using the unphosphorylated Pol II antibody (8WG16).

**Supplemental Figure S7. Boxplots show the difference in PRO-seq signal between DHS+ and DHS− TIRs**. Boxplot shows the difference in PRO-seq read counts between dREG+DHS− and dREG+DHS+ TIRs. The Y axis represents the number of reads found within 250 bp of each TIR.

**Supplemental Figure S8. Novel elements discovered using dREG frequently overlap transcription factor ChIP-seq peaks**. (A-C) Plots shows the number of TIRs discovered using dREG, but not found in DNase-I hypersensitivity or H3K27ac ChIP-seq (Y-axis) for nine PRO-seq datasets. Seven datasets were used from K562 cells (G1-8), one dataset was used from GM12878, and one from HCT116. The number of novel dREG sites overlapping transcription factor ChIP-seq peaks (dark gray),H3K4me1 ChIP-seq peaks (gray), or not overlapping either (light gray) are shown. (A) Shows the number of novel TIRs; (B) shows the number of all TIRs; (C) Shows the number of novel TIRs for a K562 dataset (G1) compared to what is expected for randomly positioned peaks that are the same size.

**Supplemental Figure S9. Overlap between dREG and GRO-cap**. (A) Venn diagram shows the overlap between GRO-cap and dREG. The intersection shows the number of GRO-cap sites. (B) Shows an example of a TIR discovered using dREG, but missed by GRO-cap.

**Supplemental Figure S10. Novel dREG TIRs overlap local increases in histone marks associated with enhancers**. Meta plots show the raw signal for H3K27ac, H3K4me1, and H3K4me3 near TIRs identified using dREG, but were not found in peak calls for H3K27ac or DNase-I hypersensitivity.

**Supplemental Figure S11. Histogram shows the distribution of jaccard distance between dREG sites in K562 cell with respect to DNase-I hypersensitivity sites in ENCODE reference cell types**. Jaccard distance was calculated for (A) all dREG sites in K562 cells, (B) dREG sites in K562 cells that do not overlap with DNase-I hypersensitive sites, and (C) dREG sites in K562 cells that do not overlap with DNase-I hypersensitive sites nor with H3K27ac ChIP-seq peaks. Major cell types among the outliers were colored and labeled.

**Supplemental Figure S12. dREG+DHS− sites do not reflect clonal differences in between K562 cells grown by ENCODE and by our lab**. Heatmaps show raw signal for two replicates of ATAC-seq data from K562 cells grown in our lab and clonally related to K562 cells used to produce PRO-seq data. Data is shown near dREG+DHS+ (n= 29,828) and dREG+DHS− (n= 7,350). Sites were ordered by dREG score.

**Supplemental Figure S13. Accuracy of classifying DHS status of dREG TIRs**. Receiver operating characteristic (ROC) plot shows the accuracy of predicting whether a TIR identified using dREG was also identified as a DHS using ENCODE DNase-I hypersensitivity data. Classification was performed using 100 transcription factor ChIP-seq datasets in K562 cells (black, auROC= 0.88) or the dREG score alone (gray, auROC= 0.75).

**Supplemental Figure S14. Pol III initiation motifs are enriched in dREG+DHS− TIRs**. Scatterplot shows the fraction of TIRs that have DNA sequence motifs representing PSE and a TATA box in the right orientation with respect to each other and with a spacing of 10-40 bp between them. Motifs were obtained from (James Faresse et al. 2012) and are shown on top. P-values comparing dREG+DHS− TIRs to dREG+ POL2A ChIP-seq+ TIRs are denoted by an asterisk (** *p* < 1e-5; * *p* < 5e-3; Fisher’s exact test).

## Supplementary Tables

**Supplemental Table S1:** Sources of PRO-seq data used in dREG model training and evaluation. Columns show the data identification number, cell line, number of mapped sequence reads, informative sites in which dREG scores were computed, reference, and the role (training or holdout validation) in this particular study.

| Data ID | Cell line | Mapped Reads | Informativ e Sites | GEO | Protocol | Reference | Role |
| --- | --- | --- | --- | --- | --- | --- | --- |
| G1 | K562 | 374,946,808 | 17,120,769 | GSM1480327 | PRO-seq | (Core et al. 2014) | Training |
| G2 | K562 | 18,129,333 | 4,189,045 | GSM1480325 | GRO-seq | (Core et al. 2014) | Training |
| G3 | K562 | 57,520,888 | 9,201,344 | GSM3452725 | PRO-seq | Herein | Training |
| G5 | K562 | 71,452,942 | 8,678,964 | GSM2361442<br>GSM2361443 | PRO-seq | (Vihervaara et al. 2017) | Training |
| G6 | K562 | 26,972,822 | 5,890,140 | GSM2545324 | PRO-seq | (Dukler et al. 2017) | Training |
| G7 | K562 | 27,046,373 | 5,839,230 | GSM2545325 | PRO-seq | (Dukler et al. 2017) | Holdout |
| G8 | K562 | 49,159,579 | 9,813,838 | GSM1622612<br>GSM1622613<br>GSM1622614<br>GSM1622615 | GRO-seq | (Niskane et al. 2015) | Holdout |
| GM | GM12878 | 105,936,649 | 9,738,488 | GSM1480326 | GRO-seq | (Core et al. 2014) | Holdout |
| GH | HCT116 | 118,728,260 | 11,660,354 | GSM1304424<br>GSM1304425 | GRO-seq | (Allen et al, 2014) | Holdout |
| CD4 | CD4 | 33,087,414 | 4,952,646 | GSM2265095 | PRO-seq | (Danko et al. 2018) | Holdout |
| MCF7 | MCF7 | 92,614,853 | 10,913,834 | GSM1014637<br>GSM1014638<br>GSM1014637<br>GSM1014638<br>GSM1014639 | GRO-seq | (Hah et al. 2011; Danko et al. 2013) | Holdout |
| H1 | HELA | 5,812,515 | 1,629,415 | GSE62046 | GRO-seq | (Andersson et al. 2014) | Holdout |
| H2 | HELA | 97,917,818 | 4,690,367 | GSM1474225 | mNET-seq | (Nojima et al. 2015) | Holdout |
| H3 | HELA | 80,739,276 | 20,385,352 | GSM1505438<br>GSM1505439 | mNET-seq | (Mayer et al. 2015) | Holdout |

